# Optical Fiber-Assisted Printing: A Platform Technology for Straightforward Photopolymer Resins Patterning and Freeform 3D Printing

**DOI:** 10.1101/2024.01.17.576081

**Authors:** Alessandro Cianciosi, Maximilian Pfeiffle, Philipp Wohlfahrt, Severin Nürnberger, Tomasz Jungst

**Author notes:** Correspondence to Tomasz Jungst.

## Abstract

Light-based 3D printing techniques represent powerful tools, enabling the precise fabrication of intricate objects with high resolution and control. An innovative addition to this set of printing techniques is Optical Fiber-Assisted Printing (OFAP) introduced in this manuscript. OFAP is a platform utilizing a LED-coupled optical fiber (LOF) which selectively crosslinks photopolymer resins. It allows on-the-fly change of parameters like light intensity and LOF velocity during fabrication, facilitating the creation of structures with progressive features and multi-material constructs layer-by-layer. An optimized formulation based on allyl-modified gelatin (gelAGE) with food dyes as photoabsorbers is introduced. Additionally, a novel gelatin-based biomaterial, alkyne-modified gelatin (gelGPE), featuring alkyne moieties, demonstrates near-visible light absorption thus fitting OFAP needs, paving the way for multifunctional hydrogels through thiol-yne click chemistry. Besides 2D patterning, OFAP is transferred to embedded 3D printing within a resin bath demonstrating the proof-of-concept as novel printing technology with potential applications in tissue engineering and biomimetic scaffold fabrication, offering rapid and precise freeform printing capabilities.

## Introduction

Vat photopolymerization (VPP) is a class of light-based additive manufacturing (AM) technologies which enables fabrication of 3D objects by light. A well-known example of these technologies is Digital-light processing (DLP). DLP printers consist of a UV light source and a digital micromirror device that selectively cure, a photo-sensitive resin with micron-level resolution and high structural complexity and generates constructs layer-by-layer^1^. However, to achieve high precision and highly resolved features, the size of the projection is restricted, and long printing times are required for the fabrication of large structures^2^. A reduction of the printing time (< 10 min for 5 mm thick constructs) has been introduced with continuous liquid interface production^3,4^. Nevertheless, this VPP fabrication technology is characterized by a costly and complex set-up and its efficiency is limited to the processing of low viscosity resins^2^. The introduction of the computed axial lithography (CAL) method, inspired by computer tomography, marks an additional significant progress for VPP in terms of printing times and variability of resin. CAL enables the fabrication of 3D objects with 0.30 mm features in a matter of seconds. Objects are achieved by rotating a synthetic photopolymer-laden reservoir within a dynamic light field. Back projections of a computer assisted digital model accumulate a light dosage, overcoming the gelation threshold and inducing material solidification exclusively in voxels corresponding to the designed 3D construct^5^. The notable potential of CAL has been demonstrated in biofabrication by volumetric bioprinting of centimeter-scale living tissue constructs in seconds using a gelatin methacryloyl (gelMA) photoresin^6^. GelMA-based hydrogels are widely used due to their compatibility with photocrosslinking strategies that preserve their shape in physiological conditions^7,8^. Nevertheless, the methacryloyl moieties of gelMA undergo chain-growth free-radical polymerization which yields to inhomogeneous polymer networks^9^. Suitable alternative resins for VPP platforms are using thiol-ene click chemistry^10^, which is a crosslinking approach that yields to highly organized hydrogel networks^11^. This polymerization strategy involves an orthogonal reaction between a carbon-carbon double bond, which is often grafted on the gelatin backbone^12^, and a thiol group of a crosslinker molecule. Furthermore, the thiol-ene reaction features a high reaction kinetic, oxygen insensitivity, compatibility with low concentration monomers, and limited hydrogel shrinking and polymerization stress^13–15^. Allyl-modified gelatin (gelAGE) is an established biomaterial platform which undergoes thiol-ene click reaction in presence of a thiolated crosslinker and a photoinitiator (PI) ^16,17^. VPP technologies often exploit resin formulations with photoabsorber (PA) to attenuate the light penetration depth and to withdraw photons from the PI, thus delaying the radical formation and yielding to more resolved features^18,19^. A common strategy for light-based 3D printing involves incorporating PA such as non-cytotoxic food dyes like tartrazine^20^ and Ponceau 4R^21^, anionic azo dye^22^, and 2-hydroxy-4-methoxy benzophenone-5-sulfonic acid^23^, into the resin. Similar to the thiol-ene click reaction of gelAGE, alkynes moieties can undergo thiol-yne click reaction in the presence of a thiolated crosslinker. Although it was used in polymer chemistry, this crosslinking strategy has been scarcely applied in the field of biofabrication with only few examples^24^.

In this study, a novel fabrication strategy named optical fiber-assisted printing (OFAP) is introduced using a benchtop and cost-effective LED-coupled optical fiber (LOF) to selectively crosslink photosensitive resins in 2D and 3D. The seamless integration of the LOF with an automated platform allows to precisely control its spatiotemporal position, irradiance, and exposure time. This enables the fabrication of photopatterned structures characterized by progressive elements (e.g., star-shaped constructs) trough on-the-fly adjustment of their fiber diameter. The developed set up is straightforward and easy to reproduce since it can be readily integrated with other automated processing technologies (e.g., robotic arms, fused deposition modeling (FDM) 3D printer), and yields micrometers-scale resolution. In addition, GelAGE-based resins are developed to meet the needs of OFAP. Tartrazine (yellow food coloring FD&C Yellow 5, E102) and Allura red AC (red food coloring FD&C Red 40, E129), whose absorbance spectra cover the UV-Vis light spectrum, are integrated with the resin precursor for the photopolymerization of structures with enhanced fiber resolution. Moreover, this work highlights the introduction of a novel gelatin-based resin functionalized with propargyl glycidyl ether (GPE) and featuring alkynes moieties. Furthermore, the alkyne moieties show absorbance properties in the near-visible UV range (405 nm) that enhanced the resolution of the photopatterned fibers without the use of PA. It is demonstrated that the great versatility of the OFAP fabrication platform permits to draw with the LOF directly within the resin bath. As proof-of-concept, the optical fiber is immersed into a gelAGE-based resin-laden vat and used for successful freeform 3D printing of centimeter scale self-standing structures within few minutes (3-5 min for 8 mm thick construct). The upscaling of OFAP from photopatterning towards 3D printing introduces a novel printing technology for VPP since, to the best of our knowledge, this work represents the first example of optical fiber-based embedded 3D printing.

## Results and discussion

### OFAP photopatterning of gelAGE-based resin with real-time adjustment of fiber resolution and multilayer fabrication

The OFAP photopatterning setup involves a x-y-z stage from a commercially available bioprinter integrated with the LOF, which perpendicularly irradiates a resin-laden vat at a specific optical fiber-to-vat distance (gap) (Fig.1a). The LOF consists of a quartz optical fiber featuring a core diameter (d) of 0.25 mm and a numerical aperture (NA) of 0.66. Trigonometry was used to determine the theoretical diameter of the light spot on the gelAGE-based photoresin surface (d_1_). The electromagnetic radiation travelling through air formed a cone with dimensions directly correlated to the NA of the optical fiber. The NA is defined as the product of the medium refractive index (air) and the sinus of the half angle of the light cone (Ѳ) (Equation 1) ^25^.

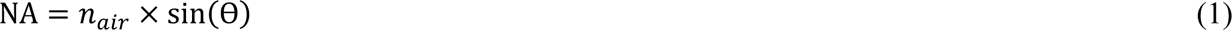

The widening of the light spot in correspondence of the resin interface (a) was and is defined as the product of the gap and the tangent of Ѳ (Equation 2).

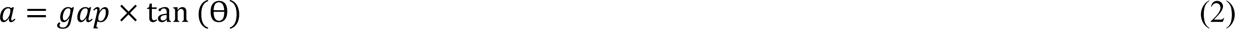

Eventually, d_1_ was calculated by using Equation 3 after introducing in the Equations 1 and 2 the features of the optical fiber (NA = 0.66 and d = 0.25 mm).

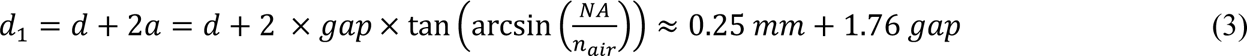

**Fig.1:**
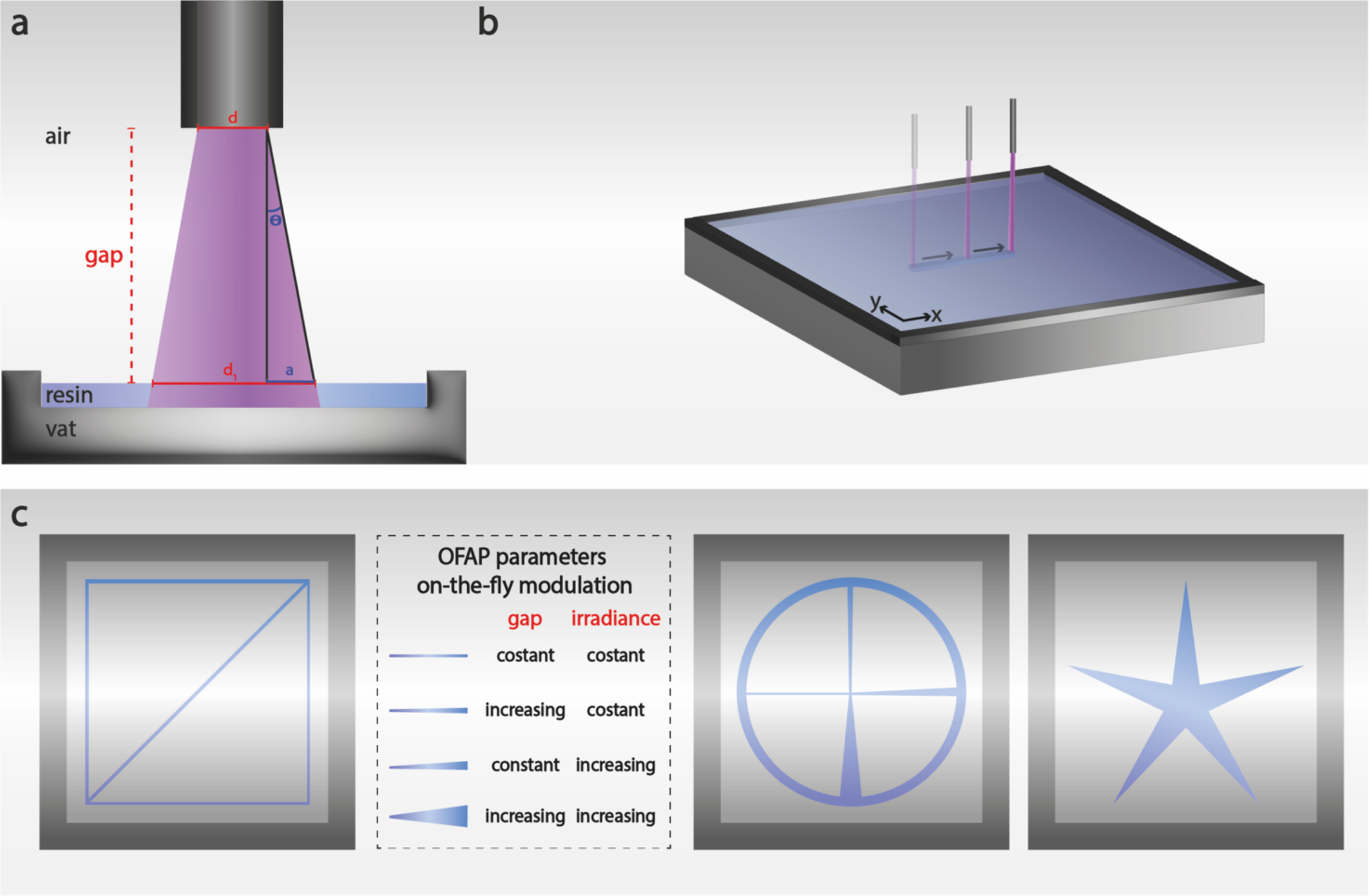
Schematic of the LOF and the OFAP photopatterning process for standard constructs and progressive elements fabrication. **a** Representation of the physical parameters of the LOF and the OFAP photopatterning process. Gap: optical fiber to vat distance; d: optical fiber inner diameter; d_1_: light cone diameter at the resin-air interface; Ѳ: sinus of the half-angle of the light cone; a: widening of the light spot, at the resin surface **b** Illustration of the automated OFAP photopatterning approach. **c** Schematic of the photopatterned structures and table describing the on-the-fly adjustments of the gap and irradiance and their influence on the fiber resolution. A 10 mm-side square design used for the fiber resolution assessment (left), a structure featuring different progressive elements and a start-shaped construct (right).

Equation 3 enabled to calculate the theoretical diameter of the light spot emitted by the OFAP platform as function of different gap distances. Particularly, gaps of 0.05, 0.15, and 0.25 mm correspond to theoretical fiber diameters of 0.34, 0.51, and 0.69 mm, respectively.

The integration of a LOF with an automated platform enabled precise spatiotemporal control for photopatterning gelatin-based photosensitive resins (Supplementary Movie 1, Fig.1b). A 10 mm-side square structure, with one diagonal, was photopatterned and used throughout the study as standard construct to assess the OFAP printing resolution of different resin formulations (Fig.1c, left). The OFAP setup allowed real-time adjustment of variables such as irradiance and gap, thus enabling the fabrication of constructs with progressive features (Fig.1c, right).

OFAP photopatterning is focused on a spatially controlled irradiation. The first step in this study was to investigate the effects of the physical parameters of the OFAP set-up on the construct resolution. A holistic approach was used to understand the interplay of the OFAP photopatterning features as gap, vat thickness and irradiance and their effects on the fiber resolution (Fig.2a). The initial hypothesis based on the theoretical light spot diameter values indicated a linear correlation between the increasing gap distances and the construct resolution as consequence of the widening of the irradiated photoresin area. Indeed, depending on the irradiance of the LED which can be adjusted between 16 and 48 mW cm^-2^ through a built-in potentiometer, the fiber diameter of the photopatterned constructs spanned from roughly 0.15 to more than 0.45 mm, while increasing the gap from 0.05 to 0.25 mm (Fig.2a). Expectedly, the measured fiber diameters were smaller compared to the theoretical ones. This phenomenon could be explained by considering the approximately gaussian distribution of the LED light spectrum^26^, which was characterized by an irradiance maximum in the center followed by a steep decrement (full width at half maximum = 15 nm) thus yielding to a photopatterned fiber featuring a fully crosslinked core and weakly cured edges. Beside the gap, which influenced the diameter of the light cone as demonstrated in Equation 3, another important physical parameter that affected the photopatterning resolution of OFAP was the irradiance. A low LED intensity resulted in a reduced energy transmitted to the gelAGE-based photoresin thus generating fewer radical species per unit of time and eventually yielding to a smaller fiber diameter^27^. The experimental data reported in Fig.2a confirmed the widening of the fiber diameter with the increase of the irradiance. Taken together, the close intercorrelation between gap and irradiance demonstrated to have a direct effect on the OFAP resolution while the vat thickness was also evaluated and showed a more relevant impact on the photopatterned construct stability than its resolution. The light emitted by the optical fiber travelled through air and was refracted by the gelAGE-based photoresin precursor. The hydrogel precursor formulation also contained a 4-arm thiolated polyethylene glycol (PEG4SH) and lithium phenyl-2,4,6-trimethylbenzoylphosphinate (LAP) which is a well-established water-soluble PI. Nevertheless, LAP is characterized by a relatively low molar absorptivity at 405 nm (ε ≈ 30 M^−1^ cm^−1^; 0.05 cm^−1^ absorptivity with 0.05 wt%)^28^. The data reported in Fig.2a showed an increased instability of the photopatterned structure while increasing the inner thickness of the vat from 0.10 to 0.20 mm with a more evident effect when the thickness reached 0.50 mm. This phenomenon could be associated with an insufficient curing depth for low irradiance values. A similar research study conducted on gelAGE using LAP confirmed a decreased curing depth using near-visible UV light while increasing the thickness of the photocrosslinked construct^29^.

**Fig.2:**
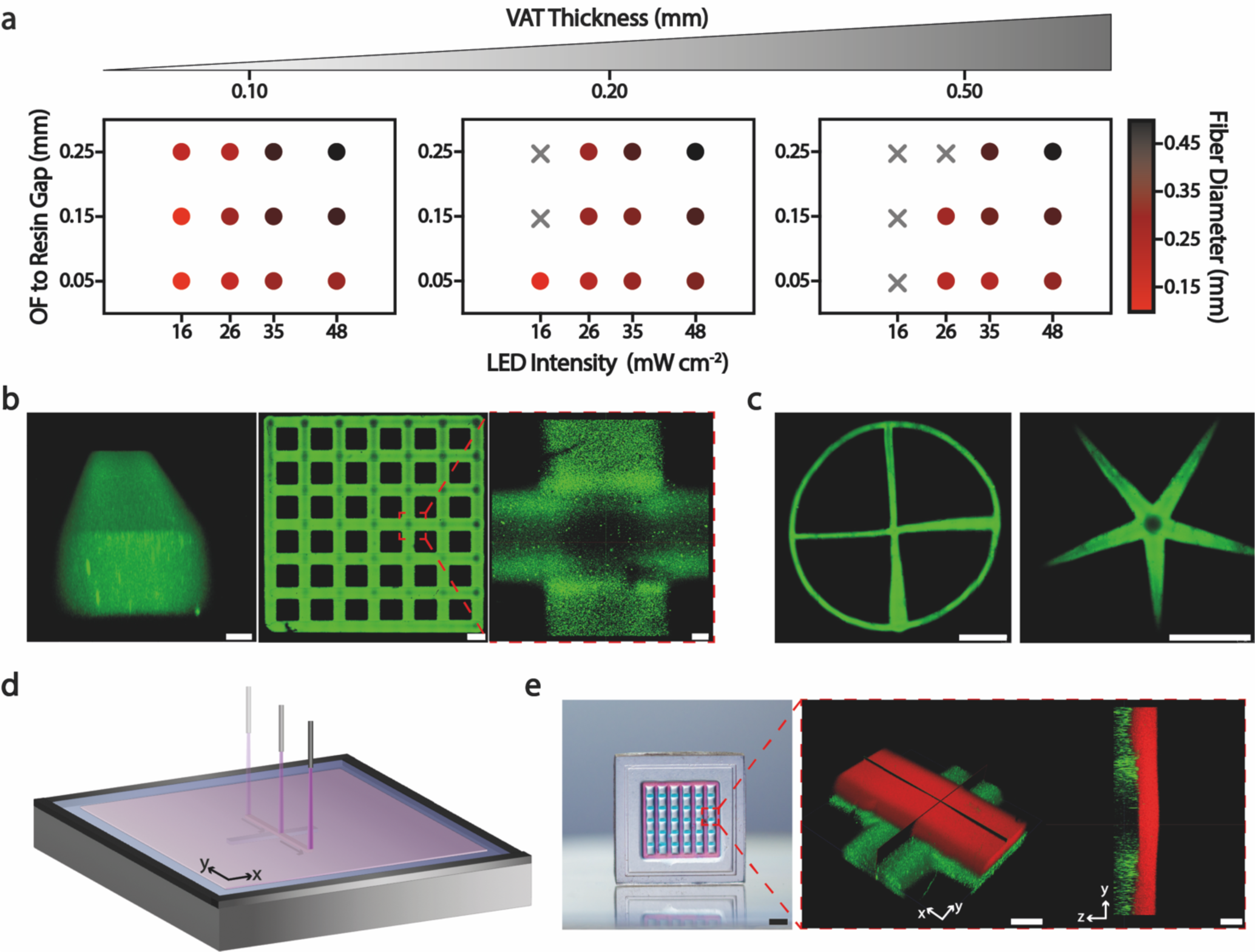
OFAP photopatterning of gelAGE-based resins. Fabrication of lattice, and multilayer structures and constructs with incremental features. **a** Interdependency of the OFAP printing process parameters (e.g., vat thickness, irradiance, gap) and their influence on the patterned fiber resolution using a gelAGE-based hydrogel precursor (5% gelAGE, 10% PEG4SH, 0.1% LAP). One sample analyzed per each point. Data are reported as mean values, n = 5 repeated measurements. **b** Photopatterning of a lattice structure with a focus on the single fiber (left) and their intersection (right), FITC-dextran was used as fluorescent dye. Scale bars (from left to right): 0.05 mm, 0.50 mm, 0.05 mm. **c** Photopatterned structure featuring incremental details (left) and a start-shaped geometry (right). Scale bars: 3 mm. **d** Schematic of the OFAP multilayer fabrication. **e** Multilayer gelAGE-based lattice structure fabricated with two adjacent layers stained with fluorescent dyes (green: FITC-dextran, red: Texas red-dextran) characterized by distinct excitation spectrum. Scale bars (from left to right): 3 mm, 0.10 mm, 0.20 mm.

A confocal image of a single gelAGE-based photopatterned fiber featuring a thickness of roughly 0.20 mm and a diameter of around 0.30 mm is depicted in Fig.2b, left. Furthermore, a single-layered lattice structure fabricated with high control, particularly on the crossover points where the irradiance overlapped, is reported (Fig.2b, right).

The understanding of the interplay between the gap and LED intensity was exploited to fabricate structures characterized by gradual features with high precision, by varying these OFAP physical parameters during the manufacturing process. Indeed, the fully automated OFAP set-up enabled to change the irradiance together with the optical fiber z position almost instantly, while patterning. The z resolution of the patterned structure is correlated to the curing depth which describes the thickness of the cured construct as function of the irradiation dosage^30^. To show the application of OFAP photopatterning for easily recreating structures with progressive elements, a design featuring four fibers perpendicularly oriented to each other and connected to the center point of the framing circle was showed (Fig.2c, left). Particularly, the fiber on the left was fabricated by simply maintaining both the gap and the irradiance (average fiber diameter around 0.48 mm) constant. The fiber on the top was patterned from the center point of the circle by gradually increasing the gap while maintaining the irradiance constant. In this case only a small incremental detail (from around 0.50 to 0.70 mm) could be noticed compared to the initial standard fiber. Nevertheless, on the fiber on the right, a larger increment was observed (from around 0.60 to 1.10 mm) when the irradiance was increased while keeping the gap constant. Eventually, the combined gradual increase of the irradiance and the gap resulted in an even more noticeable progressive fiber (from around 0.60 to 1.30 mm). Additionally, a star-shaped photopatterned structure featuring large incremental details (from around 1.40 to 0.10 mm) was reported (Fig.2c, right).

The OFAP set-up was also demonstrated to fabricate multilayer structures (Fig.2d) by exploiting the thermoreversible properties of the gelAGE. Particularly, in the multilayer fabrication approach the first gelAGE-based resin layer fluorescently stained with a fluorescein isothiocyanate (FITC)-labeled dextran was photopatterned and the remaining precursor solution was physically cured by temperature decrease (around 4 °C). The second layer of gelAGE-based resin containing fluorescent red-dextran was poured on top of the first layer and photopatterned. Eventually, the non-crosslinked precursor of both layers was rinsed to yield the final construct (Fig.2e, left). Confocal microscopy was used to highlight the lattice structure and the clear separation with absence of mixing across the two layers (Fig.2e, right).

### Photoabsorbers and triple bond gelatin-grafting to enhance the OFAP photopatterning resolution

The resolution of VPP techniques is mainly correlated to their printing hardware, although confining the photopolymerization within the irradiated voxel remains one of the challenges of these fabrication technologies^31^. Indeed, the scattering of light and the radical diffusion can influence the resin absorbance, at the irradiation wavelength, thus leading to undesired polymerization around the edges of the irradiated spot and affecting the XY plane resolution^31^. This parameter as well as the z resolution can be improved by adjusting the irradiation as already demonstrated in this study (Fig.2a), although the photoresins can be further tailored introducing PA. These are non-reactive species featuring absorbance at the irradiation wavelength thus interfering with the PI and reducing the light scattering and the overcuring^32,33^. Food dyes represents ideal PA as they are cost-effective, readily available, and cytocompatible^34^. The UV-Vis spectra of two food dyes as tartrazine (λ_max_ = 410 nm) and Allura red AC (λ_max_ = 510 nm) were measured (Fig.3a, left). These PA featured high hydrophilicity, cytocompatibility and for the tartrazine synchronized absorption wavelengths with the 405 nm LOF used for the photopatterning experiments^34,35^. Allura red AC was selected as negative control to highlight the importance of the synchronization of the PA absorbance and the irradiance wavelength^36^. The two PA were added to the gelAGE-based hydrogel precursor solutions and their effect on the photocrosslinking efficiency were assessed using photorheology and measuring the hydrogel sol fraction (mass loss at day 1) values. A time sweep analysis (Fig.3a, center) was performed on the control gelAGE (5%)-based hydrogel precursor solution featuring PEG4SH (10%) as crosslinker, and LAP (0.1%) as PI. A 0.02% of both tartrazine and Allura red AC was added to the control formulation. As expected, the trends of the storage modulus demonstrated a clear influence of both PA, particularly tartrazine, on the photocrosslinking kinetic of the gelAGE-based hydrogel formulation. Indeed, during the UV-Vis (390-500 nm) light exposure of the precursor solutions the storage modulus of the control formulation showed an almost instant (around 1 s) steep increment, whilst the presence of the PA greatly delayed the photocrosslinking reaction with a steep increment of the elastic modulus only around 5 s after light exposure. Furthermore, the highly efficient absorbance property of the tartrazine-laden precursor was remarked by a less pronounced increment, after crosslinking, of its storage modulus with invariably smaller values compared to the Allura red AC-laden hydrogel formulation. Nevertheless, it is worth mentioning that the storage modulus trend of the Allura red AC-laden hydrogel precursor was influenced by the technical constrains of the photorheological set up, which was used to best approximate the OFAP photopatterning conditions. Indeed, this installment consisted of a UV-Vis lamp equipped with a filter narrowing the electromagnetic spectrum to a range spanning from 390 to 500 nm. Theoretically, using the LOF, the time sweep trend of the Allura red AC-laden precursor should be similar to the gelAGE-based control solution, due to the lack of synchronization of this PA absorption wavelength (α_max_ = 510 nm) with the 405 nm LED. The sol fraction analysis conducted with the LOF confirmed this hypothesis. This analysis was performed to quantitively address the photocrosslinking efficacy of the three gelAGE-based hydrogel precursor solutions (Fig.3a, right). The sol fraction is a quantitative and straightforward analysis of the photocrosslinking efficacy and defines the fraction of non-crosslinked polymer washed out of the hydrogel network after 24 h^37^. The control hydrogel formulation showed a sol fraction (4.3 ± 3.9%) significantly smaller than the gelAGE-based hydrogel formulation containing the tartrazine (39.0 ± 3.2%) confirming its high absorbance and effect on the hydrogel photocrosslinking efficacy. Furthermore, the lack of absorbance wavelength synchronism between the LOF and the Allura red AC and thus its poor effect on the photocrosslinking reaction was confirmed by its sol fraction value (1.1 ± 5.3%), which was comparable to the control precursor solutions.

**Fig.3:**
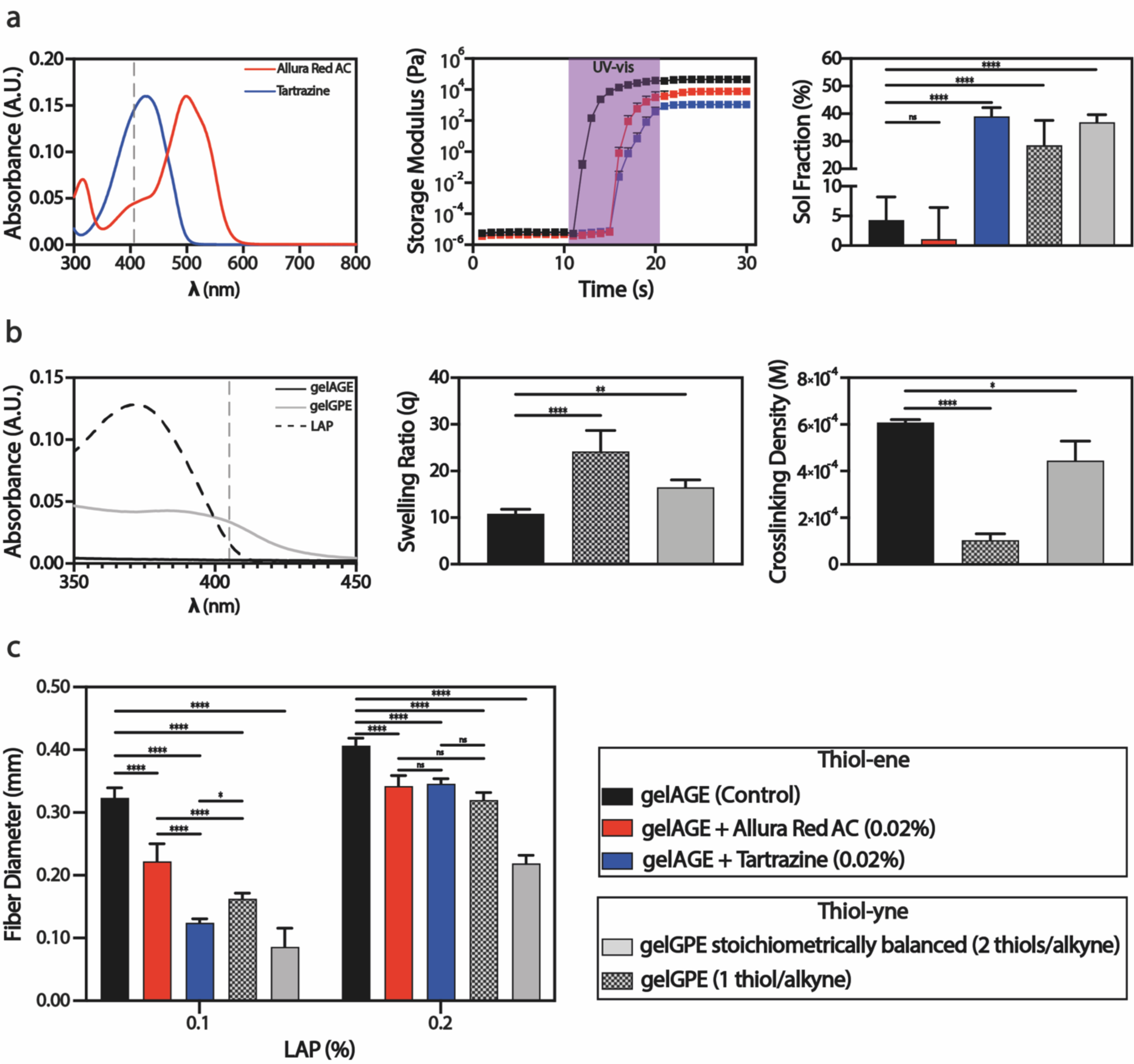
Quantitative evaluation of the effect of photoabsorbers and alkyne moieties on the photopatterned fiber resolution. **a** Introducing PA, Allura red AC and tartrazine, into the gelAGE-based precursor formulation. UV-Vis spectrum of both PA (0.002%, PBS as solvent, left). Photo-rheological oscillatory analysis: Time sweeps trends of the gelAGE-based precursor without (control) and with Allura red AC and tartrazine (center). Data are reported as mean values ± SDs, n = 3 independent experiments. Crosslinking efficacy expressed as sol fraction (mass loss after 24 h) of the gelAGE-based hydrogel (control) compared to the gelAGE-based precursors containing Allura red AC and tartrazine, respectively, and the gelGPE-based hydrogels both stoichiometrically balanced and in thiol defect (right). Data are reported as mean ± SDs, n = 6 independent experiments. Statistical significances are determined based on a one-way ANOVA. **b** Absorbance properties of the gelatin featuring alkyne moieties and thiol-yne click chemistry to tailor the hydrogel crosslinking density. UV-Vis spectrum of gelGPE in comparison to gelAGE and LAP (5% of gelAGE and gelGPE, 0.1% of LAP, PBS as solvent, left). Swelling trends (after 24 h) of the gelAGE-based hydrogels in comparison to the gelGPE-based hydrogels both in stochiometric balance and in thiol deficit (center). Data are reported as mean ± SDs, n = 6 independent experiments. Statistical significances are determined based on a one-way ANOVA. Crosslinking density trends of the gelAGE-based hydrogels compared to the gelGPE-based hydrogels (right). Data are reported as mean ± SDs, n = 3 independent experiments. Statistical significances are determined based on a one-way ANOVA. **c** Quantitative evaluation of the photopatterned fiber resolution of gelAGE-based hydrogels (w/o PA) and gelGPE-based hydrogels (stoichiometrically balanced and in thiol deficit), as function of the LAP concentration. Data are reported as mean ± SDs, n = 5 repeated measurements per each group (Supplementary Fig.2). Statistical significances are determined based on a two-way ANOVA. **a-c** The significance of p is indicated by * = p < 0.05, ** = p < 0.01, *** = p < 0.001, **** = p < 0.0001.

Hydrogels are generally transparent in the near-visible UV and visible range, thus PA are useful tools for controlling the photocrosslinking^31^. Nevertheless, in this study we introduced a gelatin-based hydrogel featuring alkyne groups (gelGPE) which was characterized by light absorbance properties at 405 nm thus representing an ideal alternative to PA for OFAP photopatterning (Fig.3b, left). The photorheological analysis demonstrated the effect of the alkyne moieties on the crosslinking, as indicated by the delayed steep increment of the storage modulus of the gelGPE-based formulations compared to the gelAGE-based control (Supplementary Fig.1). Furthermore, alkyne moieties can undergo a thiol-yne click chemistry crosslinking reaction in the presence of a thiolated crosslinker, and a PI. The terminal alkyne moiety, contrary to the monofunctional vinyl group involved in the thiol-ene reaction, is bifunctional and can be used to further tailor the intrinsically higher hydrogel crosslinking density, by adjusting the stochiometric ratio between carbon-carbon triple bond and the thiol moieties^38^. The photopolymerization of the gelGPE-based hydrogel precursor solution involved the addition of a thiyl radical to an alkyne moiety, generating a carbon-centered radical that abstracted a hydrogen from another thiol thus yielded to a vinyl sulfide group. At this stage, differently from the thiol-ene reaction, the vinyl sulfide group reacted with another thiyl radical eventually yielding to a dithioether^39^. The thiol-yne reaction yields highly crosslinked hydrogel networks similarly to a multifunctional chain-growth polymerization whilst keeping a step-growth mechanism which often results in a slightly delayed gel point and higher network homogeneity^38^. After complete thiol-yne reaction, each functional group of the gelGPE was combined with two thiol moieties thus establishing the alkynes as difunctional, instead of the monofunctional allyl groups involved in the thiol-ene. Indeed, the thiol-yne click reaction enabled to obtain multifunctional and denser polymeric networks^40^. The crosslinking density (π_x_) of the gelAGE- and gelGPE-based hydrogel networks was estimated using the theory of rubber elasticity^41^ (Equation 4) assuming the materials are incompressible (Poisson’s ratio (ϖ) = 0.5), since hydrogels are characterized by extremely high-water content and show large strain under low loading conditions^42^.

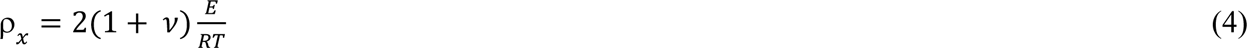

The π_x_ was calculated from Equation 4 where R is the gas constant, T the temperature, and E the elastic modulus measured with uniaxial unconfined compression tests on the gelAGE- and gelGPE-based hydrogels. The gelGPE-based hydrogel precursor solutions were formulated by using a stoichiometric balance of thiol and alkynes moieties (2 thiols/ alkyne) and in thiol deficit (1 thiol/ alkyne). In both precursor solutions a 5% concentration of gelGPE with either 2.6 or 5.2% of PEG4SH, and 0.1% of LAP was used. The investigation of different concentrations of thiols in the gelGPE-based hydrogel formulations was designed to better highlight their effect on the OFAP photopatterning. Indeed, the kinetic of the thiol-yne reaction is rather correlated to the thiol instead of the alkyne concentration since the chain transfer is the rate determining step^38^. The estimated crosslinking density value of the gelGPE-based hydrogel featuring a thiol deficit was 1.03 ± 0.26 ×10^-4^ M. This was, expectedly, four times smaller compared to 4.40 ± 0.82 ×10^-4^ M of the gelGPE-based hydrogels with stoichiometrically balanced alkynes and thiols. The gelAGE-based hydrogel showed a crosslinking density of 6.06 ± 0.11 ×10^-4^ M which was significantly higher than both gelGPE-based hydrogels (Fig.3b, right). This phenomenon confirmed the high absorbance properties of the gelGPE which theoretically should have yielded to a denser polymer network than gelAGE, due the nature of the thiol-yne crosslinking mechanism.

A further confirmation of this hypothesis was shown by the sol fraction values (Fig.3a, right) of the gelGPE-based hydrogel stoichiometrically balanced (36.9 ± 2.7%) and in thiol deficit (28.5 ± 9.1%). These values are indeed in line with those of the tartrazine which already showed to have a synchronized absorption spectrum, as the gelGPE, with the LED of the OFAP set-up. Moreover, π_x_ influences other physicochemical properties, beside the mechanical stiffness, as the solute diffusivity^43^. Indeed, the swelling trend (Fig.3b, center) of the gelGPE-based hydrogel in thiol deficit which was characterized by a low crosslinking density showed significantly higher swelling (24.1 ± 4.5%), after 24 h, compared to its stoichiometrically balanced analog (16.5 ± 1.5%) and the gelAGE-based control (10.8 ± 1.0%).

The OFAP photopatterning resolution for the different screened resins was evaluated and trends were reported (Fig.3c). The resolution of VPP processing technologies can be influenced by physical and chemical parameters as light scattering and radical diffusion, respectively. This latter parameter is strictly correlated with the crosslinking efficacy and density, although one of the challenges of VPP techniques is the precise confinement of the photocrosslinking reaction in predetermined areas of the resin^44^. Indeed, another important factor for the OFAP photopatterning process was the introduction in the resin formulations of PA and alkyne moieties (gelGPE) which limited the overgrowth (light scattering) by absorbing part of the light and challenged the PI. The fiber resolution assessment of the photopatterned resin containing 0.1% LAP, showed significantly more resolved fiber for the gelAGE-based hydrogels featuring PA compared to the control (0.323 ± 0.015 mm). Tartrazine which was characterized by a synchronized absorbance spectrum at 405 nm showed fibers with smaller diameters (0.124 ± 0.007 mm) compared to the effect of the Allura red AC on the gelAGE-based hydrogel resolution (0.222 ± 0.028 mm). Being characterized by the presence of alkynes groups which showed absorbance at 405 nm, the gelGPE-based hydrogels were expected to yield more resolved fibers compared to the control. Indeed, the stoichiometrically balanced gelGPE-based hydrogel featured fiber diameters in the order of the tartrazine-based analogs (0.086 ± 0.030 mm). Furthermore, the gelGPE-based hydrogel crosslinked in thiol deficit showed larger fiber (0.162 ± 0.009 mm) as expected since the kinetic of the thiol-yne reaction is strongly correlated to the thiol concentration. The resolution trends of the screened photoresins were invariably higher when the LAP concentration was increased to 0.2%. Nevertheless, similarly to the hydrogels featuring 0.1% LAP, these trends showed a better resolution for the resins containing PA and gelGPE compared to the gelAGE-based hydrogel control. Contrary to the resins featuring 0.1% LAP, a non-significant improvement of the resolution from the presence of tartrazine compared to the Allura red AC was noticed, similarly for the gelGPE-based hydrogel in thiol defect. This phenomenon could be explained by the double increment of PI which overcame the absorbance of the PA and of the alkyne moieties of the gelGPE.

### OFAP for the embedded 3D printing of self-standing structures

A pivotal aspect of this study was the exploitation of the versatility of the OFAP printing setup towards the 3D freeform fabrication of self-standing structures. In a proof-of-concept approach, the LOF was used to directly print 3D structures within a tartrazine-laden gelAGE-based resin bath (Supplementary Movie 2, Fig.4a). The OFAP 3D printing process was based on the controlled spatiotemporal position (feed rate = 1 or 2 mm s^-1^) of the tip of the optical fiber in direct contact with the photosensitive resin bath to draw consecutive horizontal, vertical, and diagonal patterns in the 3D space. GelAGE was already introduced as resin for stereolithography 3D printing and was characterized by a concentration around 10 to 20%^45^, thus for the OFAP 3D fabrication the gelAGE concentration was increased from 5 to 10%. The gelAGE-based resin formulation, along an increment of its concentration, was characterized by a higher LAP content (0.3%). To balance the relatively low 3D printing feed rate, the PI concentration was increased to enhance the radical production per time. Furthermore, the optical fiber used for the 3D printing fabrication featured a different core diameter and numerical aperture (d = 0.50 mm, NA = 0.63), and was made of polymer instead of quartz. The new material was ideal for the 3D printing approach since the optical fiber is in direct contact with the resin while moving, thus the more brittle and fragile quartz fiber could have been subjected to damage or even breakage. The new polymer optical fiber was characterized by different light intensity outputs spanning, from lowest to highest, from 17 to 130 mW cm^-2^ (well above the maximum intensity obtained with the quartz fiber = 48 mW cm^-2^). The adjustment of the precursor and PI concentration helped to enhance the reactivity of the material during the printing process and the LED irradiance of the new optical fiber was increased to achieve a higher crosslinking rate. The initial focus while developing this new optical fiber-based 3D printing technique was centered on the fiber tip-to-resin interface where a small buildup of cured resin was present. These small parts of cured resin were continuously detached from the optical fiber tip during the movement within the bath, thus yielding to fibers featuring slightly bumpy textures. The printing of vertical features was the most straightforward aspect of the 3D fabrication process. Indeed, while driving the optical fiber upwards (feed rate = 2 mm s^-1^) within the resin-laden bath several physical forces were involved particularly at the fiber tip-to-resin interface and helped the continuous removal of cured buildups from the fiber tip. The upward controlled movement of the fiber combined with the buoyancy of the gelAGE-based resin reservoir facilitated not only the stability of the printed construct but also the detachment of the buildups of crosslinked resin. Moreover, the localized increment of the refractive index in correspondence of the cured vertical fiber acted as optical constrain, thus favoring the propagation of the patterned front through the resin volume^46^. For the fabrication of horizontal fibers, the feed rate was decreased to 1 mm s^-1^ while keeping the LED irradiance constant. Furthermore, the light penetration depth played an important role and required further optimization. This aspect was particularly important when crosslinking horizontal lines, considering the self-standing nature of the printing constructs. The presence of tartrazine in the precursor solution was ideal to limit the light penetration depth and increase the z resolution of the structures limiting the crosslinking to the desired layer thickness.

Once the optimization of the OFAP 3D printing system was completed on both the vertical and horizontal motions, more complex 3D structures were printed (Fig.4b, right). A cube featuring an 8 mm-side with a fiber resolution around 1.7 mm was fabricated. In this structure the main aspects to optimize were the transition movements from vertical to horizontal paths which resulted in a slightly less resolved edge compared to the main fiber. Diagonal paths were introduced in an 8 mm-side square-based pyramid which featured fibers with a resolution of around 1.7 mm and a height around 8 mm. To demonstrate the flexibility of structures design of the OFAP 3D printing platform more complex analogs were fabricated (Fig.4b, left and center).

**Fig.4:**
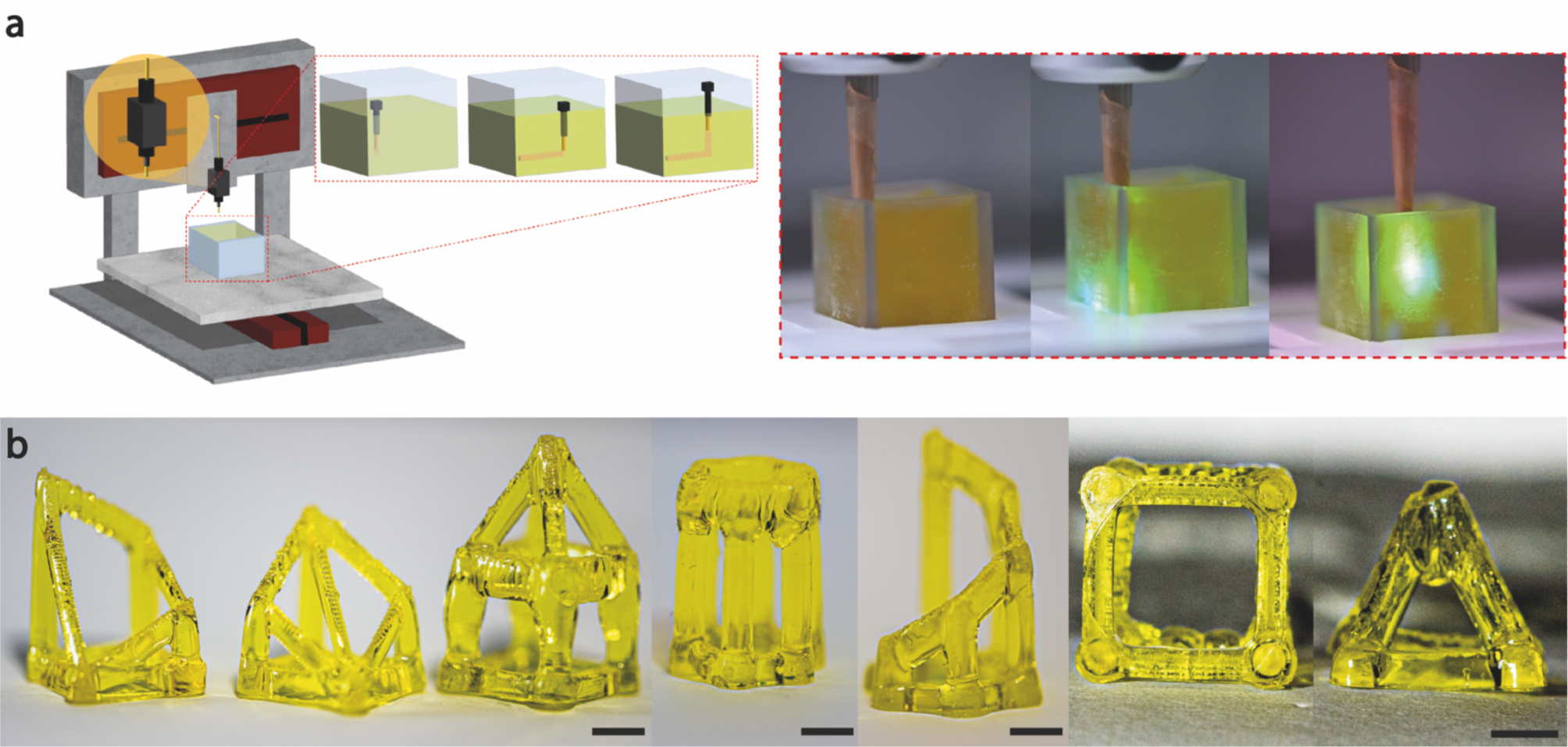
OFAP 3D fabrication of gelAGE-based constructs. **a** Schematic (left) and actual representation (right) of the 3D printing process using the OFAP platform. The LOF is immersed into the photosensitive-laden vat and its spatiotemporal position its automatically controlled to generate a 3D structure. **b** Proof-of-concept fabrication of self-standing 3D structures (e.g., hollow cube and pyramid) based on gelAGE-hydrogels with tartrazine as photoabsorber. Scale bars: 3 mm.

Taken together, the proof-of-concept 3D fabrication based on the OFAP platform showed promising results although highlighting some aspects to further optimize to enhance the quality of the prints.

## Conclusions

This study introduced a novel AM fabrication strategy, OFAP, for the precise patterning of gelatin-based photosensitive resins using a benchtop printer and a cost-effective LOF. The developed OFAP setup, integrated with an automated platform and open-source coding software (Python), allowed for precise control over the spatiotemporal position, irradiance, and exposure time of the LED light source. The holistic investigation of the effects of the OFAP parameters such as gap, vat thickness, and irradiance on the photopatterned fiber resolution demonstrated a close intercorrelation between these variables. Gap and irradiance were varied on-the-fly to adjust the printing resolution, thus fabricating constructs characterized by progressive elements. The use of tartrazine and Allura red AC as PA in the gelAGE-based resin precursor proved effective in enhancing the photopatterning resolution. Furthermore, the introduction of the novel gelGPE featuring alkyne moieties exhibited near-visible UV absorbance properties offering an alternative approach to traditional PA. The versatility of the OFAP platform was demonstrated by its successful transition from photopatterning to freeform 3D printing within a gelAGE-based resin bath. The ability to directly print self-standing structures (e.g., hollow cube and squared-based pyramid) within a low viscosity resin bath, to the best of our knowledge, represents the first example of optical fiber-based embedded 3D printing. The proof-of-concept study provided promising results and underscored the flexibility of the OFAP fabrication platform. Nevertheless, to further advance the OFAP 3D platform some improvements can be foreseen as a more engineered optical fiber tip with, for instance, hydrophobic coatings which would prevent the formation of buildups and smoothen the texture of the printed structures. Additionally, an optical fiber featuring a smaller core diameter will enhance the resolution further. The optimization of the coding scripts (Python and G-code) and the printing parameters (e.g., feed rate) could also lead to the introduction of round features in the printed constructs.

Overall, the OFAP approach offered a facile, reproducible, and versatile solution for photopatterning suggesting the use of OFAP as straightforward platform to adapt real-time the fiber resolution and generate incremental details within structures. Furthermore, OFAP freeform 3D printing of gelatin-based structures showed potential applications in various fields, including tissue engineering and biomimetic scaffold fabrication.

## Methods

### Synthesis of functionalized gelatin with allyl moieties (gelAGE)

The gelAGE synthesis was adapted and optimized based on previously published protocols^45^. Type A gelatin from porcine skin (Sigma Aldrich) was dissolved in deionized water (10%) for 2h at 37 °C. Two types of gelAGE were synthetized, G_1MM_ and G_2LH_, with only the latter featuring thermoreversibility. The nomenclature refers to the reaction conditions and the educts molar concentrations as already thoroughly indicated in our previous study^17^. Sodium hydroxide (NaOH, 2 M, Sigma Aldrich) in a concentration 0.4 or 2.0 mmol gram^-1^ of gelatin, and allyl-glycidyl ether (AGE, Sigma Aldrich) in a concentration of 12 or 24 mmol gram^-1^ of gelatin were added to the gelatin at 65 °C for 1 or 2h. The deprotonated amino groups on the gelatin backbone were linked to the less substitute carbon of the heterocyclic ring of AGE through a nucleophilic substitution. The synthesized gelAGE products were dialyzed (Spectra/Por 6, Fischer Scientific; molecular weight cut-off = 1 kDa) against deionized water and then lyophilized.

### Synthesis of functionalized gelatin with alkyne moieties (gelGPE)

The grafting of the alkyne functional groups on the gelatin backbone was inspired by the gelAGE reaction mechanism. Similarly, type A gelatin from porcine skin (Sigma Aldrich) was dissolved in deionized water (10%) for 2h at 37 °C. NaOH (2 M, Sigma Aldrich) in a concentration 0.4 mmol gram^-1^ of gelatin, and glycidyl-propargyl ether (GPE, Sigma Aldrich) in a concentration of 24 mmol gram^-1^ of gelatin were added to the gelatin at 65 °C for 2h. NaOH induced the alkaline conditions which led to the same nucleophilic substitution which caused the deprotonated amino groups on the gelatin backbone to link to the exposed carbon site on the heterocyclic GPE ring. The synthesized gelGPE polymer was dialyzed (Spectra/Por 6, Fischer Scientific; molecular weight cut-off = 1 kDa) against deionized water and then lyophilized.

### Molecular characterization of the modified gelatin-based biopolymers

Aqueous gel permeation chromatography (GPC) analysis (0.1 M NaNO_3_, 0.02 wt.% NaN_3_) of gelAGE and gelGPE was conducted using a Malvern system (Malvern) (Supplementary Fig.3). The apparatus consisted of a Viskotek GPCmax (in-line degasser, 2 piston-pump and autosampler), column oven (35 °C), refractive index (Viskotek) and SEC-MALS 20 (Viskotek) detector. Column set was 2 x A6000M (length: 300 mm, width: 8 mm, porous polyhydroxy methacrylate polymer, particle size A6000M = 13 µm). The flow rate was 0.7 mL min^-1^. Calibration was performed using polyethylene glycol standards (Malvern). The molecular weight of the PEG standards was comprised between 0.4 to 700 kDa. Samples were allowed to dissolve overnight (3 mg/mL) and filtered through a syringe filter (Thermo Fischer Scientific; 0.45 µm regenerated cellulose) prior to analysis. Proton nuclear magnetic resonance (^1^H-NMR) spectra of gelAGE and gelGPE were recorded on a Bruker Biospin 400 MHz spectrometer (Bruker) with deuterium hydroxide (D_2_O) as solvent (Supplementary Fig.4). As internal reference the signal at δ = 4.79 ppm was used. The phenylalanine peaks at δ = 7.45 - 7.25 ppm were used as reference signals for AGE and GPE modifications and calibrated to 5 protons. To determine the degree of modification (DoM) of both functionalized gelatins, the proton integral of the allyl and alkyne moieties was compared to the phenylalanine (Phe). On average, 1 gram of gelatin contains 21 milligrams of phenylalanine (2.1%). The composition of acid-treated porcine skin gelatin was acquired from the gelatin handbook of the gelatin manufacturers institute of America (GMIA). The DoM is calculate based on Equations 5, 6, and 7.

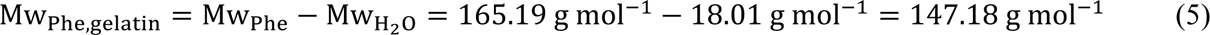

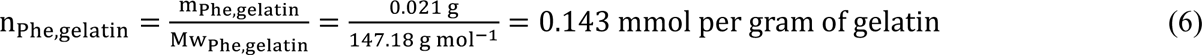

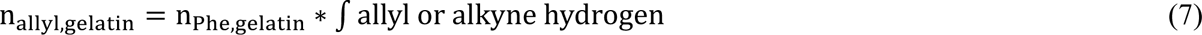

### Photoresin precursor solutions preparation

The freeze-dried gelAGE (G_1MM_) and gelGPE together with a commercially available PEG4SH (JenKem Technologies), in a powder state, were dissolved in phosphate buffer saline (PBS, 1-fold, Sigma Aldrich). Eventually, the photoinitiator LAP (Sigma Aldrich) was diluted into the resin precursor (0.1 or 0.2%) from a freshly prepared stock solution (1%) in PBS. The resin precursor solutions were allowed to thoroughly dissolve at 37 °C for few minutes before the transfer in a custom-made vat. Furthermore, the photoabsorbers tartrazine (Sigma Aldrich) and Allura red AC (Sigma Aldrich) were diluted into the gelAGE-based precursor (0.02%) from a freshly prepared stock solution (0.5%) in PBS. The gelAGE (G_2LH_) featuring thermoreversibility was used only for the fabrication of the multilayer structure exploiting its physicochemical property.

### Physicochemical characterization of gelAGE-based hydrogels

The swelling and mass loss analysis of both gelAGE- and gelGPE-based photoresins was adapted from previous studies^47–49^. In brief, cylindrical hydrogels (diameter: 6 mm, height: 1 mm) were fabricated. Samples were weighed after the crosslinking to obtain the initial mass (m_initial_, _t0_) and then lyophilized to get the dry weight (m_dry_, _t0_). The other samples (n = 6) were incubated at 37 °C in PBS for 24h. Samples were blotted dry and weighed to obtain the swollen mass (m_swollen_, _t_) and lyophilized to get the dry weight (m_dry_, _t_).

The swelling ratio (q) of the hydrogels was calculated based on Equation 8.

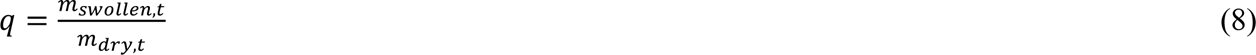

The percentage of sol fraction which corresponds to the amount of non-crosslinked polymer that leaches out form the hydrogel network after 24h, was calculated based on Equation 9.

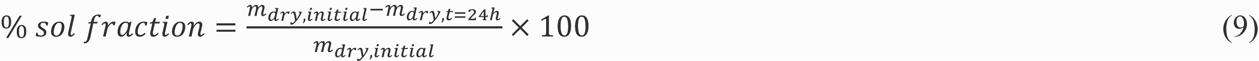

Thereby, m_dry_, _initial_ was calculated using the actual macromer fraction as reported in Equation 10 and 11.

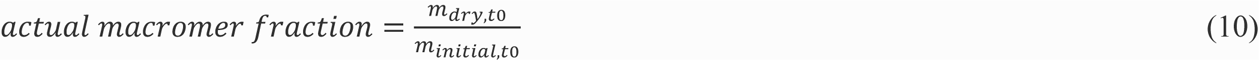

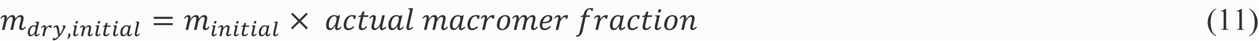

### Mechanical properties and crosslinking density measurement

The mechanical properties of the hydrogels were measured in an unconfined, uniaxial compression test, using a dynamic mechanical analyzer (BOSE, ElectroForce 5500 system). Cylindrical samples after crosslinking (diameter: 5 mm, height: 3.5 mm, n = 3) were subjected to a strain ramp until 25% compression of the sample height was reached (0.01 mm s^-1^, preload buffer time 20 s). The Young’s modulus (E) was calculated from the slope of the stress-strain curve. E was used for the estimation of the sample crosslinking density using the theory of rubber elasticity (Equation 12) assuming that the material was incompressible (ϖ = ½)^38^.

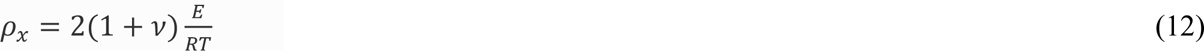

### UV-Vis spectroscopy and LED irradiance measurement

The UV-Vis analysis of both functionalized gelatin biomaterials and the photoabsobers was conducted with a spectrometer (Genesys 10S UV-Vis, Thermo Scientific). The samples were dissolved in distillated water and loaded on a transparent glass vial. Firstly, a blank was obtained using only distilled water (solvent) and subsequently each sample was measured within a wavelength spectrum spanning from 200 to 800 nm. A radiometer (Radiometer RMD, Opsytec Dr. Gröbel) was used to measure the irradiance. The optical fiber was placed at 0.15 mm from the measuring sensor and a live monitoring of the light intensity was performed.

### Confocal microscopy

gelAGE-based hydrogel precursor solutions were stained with fluorescent (FITC; MW: 500 kDa; Sigma Aldrich and Texas red; MW: 70 kDa; Thermo Fischer) dyes-labeled high molecular weight dextran. For imaging the scaffolds were placed in a Petri dish with a glass bottom slide (FluoroDish Cell Culture Dish, World Precision Instruments, FD35-100) containing few drops of water. The imaging was conducted using a confocal reflection microscope (Leica LSM SP8, Leica Microsystem) with a laser wavelength of 496 and 561 nm, respectively.

### Rheological measurements

Rheological properties of the different resin formulations were characterized using an Anton Paar MCR 702 rheometer (Anton Paar) with a 25 mm parallel plate geometry at a 500 μm gap. A solvent trap was used to limit evaporation phenomenon during the experiments. All measurements were conducted by loading the freshly prepared hydrogel precursors on the lower plate at room temperature (21 °C). Photorheological measurements (time sweep) were performed by coupling the rheometer with a UV-Vis light source (Dr. Hönle, bluepoint 4) equipped with a flexible light guide. The wavelength spectrum was narrowed using an optical filter between 390 and 500 nm. The glass of the lower enabled the UV-Vis light to crosslink the precursors using an exposure time of 10 s at 7.5 cm distance (light probe to the lower plate; light dosage = 254 mJ cm^-2^).

### Vat fabrication

The vats were designed with Autodesk Fusion 360 software and exported as .stl files. They were further processed into .sl1s files using the Prusa Slice software. Afterwards, the vats were printed using the Prusa Sl1S Speed DLP-printer. Post fabrication, the vats were cleaned and post-cured both for 5 min exposure in a UV chamber CW1S and with 2000 flashes in a light curing unit (Otoflash G171, NK-Optik). The vat dimensions were tailored depending on the type of fabrication process. The photopatterning experiments were conducted in a vat with a 21.25 mm side, an overall thickness of 5 mm and an inners thickness of 0.20 mm able to contain 60 μm of photoresin precursor solution. The 3D printing was conducted instead in larger vat with a side of 18 mm and a thickness of 20 mm enabling the printing of relatively large 3D structures.

### Setup for OFAP photopatterning and 3D printing

A benchtop and compact (56 x 106 x 166 mm) continuous and high-power output LED-light source (Silver LED, 405 nm, Prizmatix) connected with a quartz-based optical fiber featuring a 0.25 mm diameter and a NA of 0.66 was used for the photopatterning of click resins. The optical fiber was placed onto a 3D printer (3DDiscovery, RegenHU) printhead. The LOF was controlled via the Prizmatix Multi LED Ctrl software and connected to a computer, as it was the 3D printer which was controlled via the integrated 3D Discovery software. Python was used to connect and synchronize both software. In this way the whole printing process was programmed and controlled using phyton. To ensure an ideal positioning of the vat containing the resins precursor solution an FDM-printed holder was mounted on the collector plater of the 3D printer. The feed-rate was used at 1 and 2 mm s^-1^. For the OFAP 3D printing a different optical fiber made of polymer and featuring a diameter of 0.50 mm and a NA of 0.63 mm was used in the aforementioned setup, although using a different vat.

### Multilayer photopatterning

The thermoreversibility properties of the G_2LH_ type of gelAGE were used to fabricate a two-layer structure. A first layer of G_2LH_ was evenly spread onto the surface of a vat with a 0.50 mm inner thickness, then physically gelled by incubating the vat into a custom-made cooling chamber for a few minutes. The first layer was then photopatterned and cooled down to gel the unreacted resin again. Subsequently, another layer of G_2LH_ was distributed onto the first material surface and then photopatterned. Eventually, the construct was thoroughly rinsed with distilled water to remove the excess of unreacted resin.

### Fiber resolution assessment

Images of the photopatterned structures designed for the fiber resolution assessment were taken with a stereomicroscope (Leica DMS1000) using constant lighting conditions and focus. Each image was processed using the open-source software fiji. Briefly, the images were cropped and duplicated it. A threshold (Red-Yen) was applied on the original images which was subsequently eroded ad dilated. Eventually, the image calculator feature was used to sum the two images and obtain a smooth and clear image to accurately calculate the fiber diameter in different spots of the designed photopatterned structure (Supplementary Fig.4).

### Statistical analysis

All data were expressed as means ± standard deviations (SDs) for n ≥ 3. The normal distribution of the data was checked with a Shapiro-Wilk test. GraphPad Prism software (GraphPad software) was used to perform a one- or two-way analysis of variance (ANOVA) with a Dunnett’s or a Tukey’s post hoc multiple comparison test, respectively, to determine statistical significance. Values of *p* < 0.05 were considered statistically significant. * = statistically significant differences (* = p < 0.05, ** = p < 0.01, *** = p < 0.001, **** = p < 0.0001).

## Data availability

Data are available from the corresponding author upon reasonable request.

## Supporting information

Description of supplementary information files

## Acknowledgments

T.J. acknowledges the funding from the Deutsche Forschungsgemeinschaft (DFG, German Research Foundation), Project number 326998133, TRR 225 (subprojects B09 and C02). T.J. acknowledges the European Union for funding via the European Union’s Horizon 2020 research and innovation program under Grant Agreement No. 874827 (project BRAV3).

## Author contribution

A.C., T.J. designed research; A.C., M.P., P.W. S.N performed research; A.C. analyzed data; A.C. wrote the paper.

