## Supplementary information for "Optical Fiber-Assisted Printing: A Platform Technology for Straightforward Photopolymer Resins Patterning and Freeform 3D Printing"

**This pdf file includes:**

Supplementary Figures 1 to 4

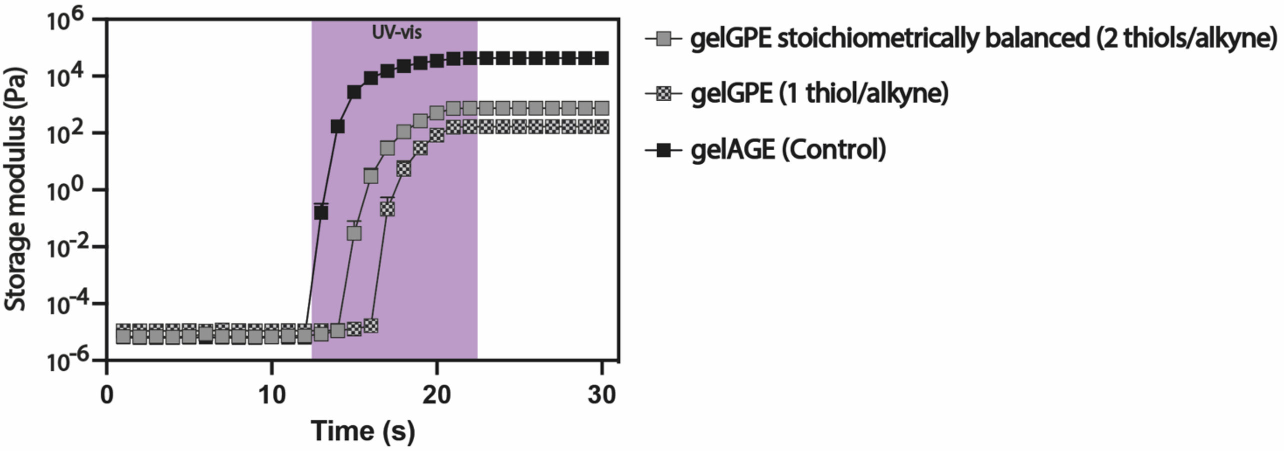


**Supplementary Fig.1: Photorheological analysis of gelGPE-based precursor solutions.** Photo-rheological oscillatory analysis (n = 3, 10 rad $s^{-1}$ oscillation frequency, 10% shear strain): Time sweeps trends of the gelAGE- (control) and gelGPE-based precursor with different stoichiometric balance (1 and 2 thiols/alkyne).

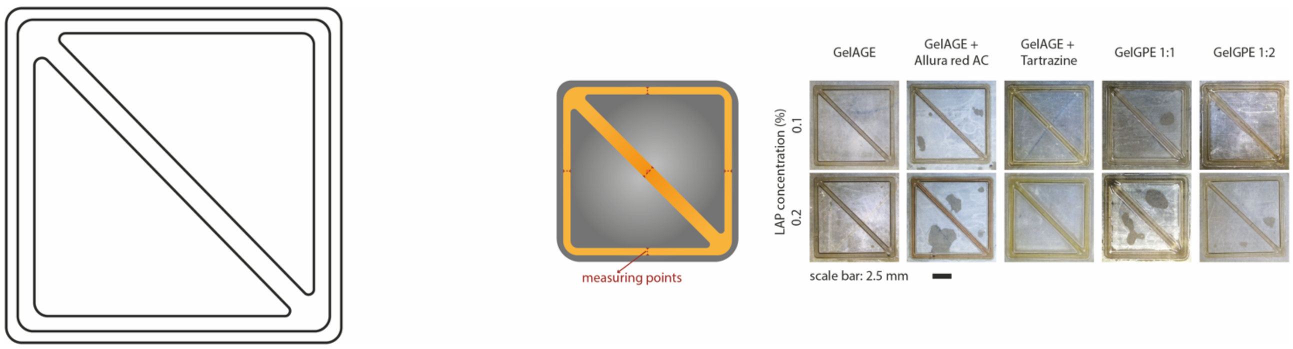


**Supplementary Fig.2: Fiber resolution assessment of different photopatterned resins.** Schematic of the designed pattern for the determination of the fiber diameter (left). Schematic of the selected measuring areas (n = 5) within the photopatterned structures along with the original stereomicroscope images of different photosensitive resins (e.g., gelAGE w/o PA and gelGPE with different molar ratios).

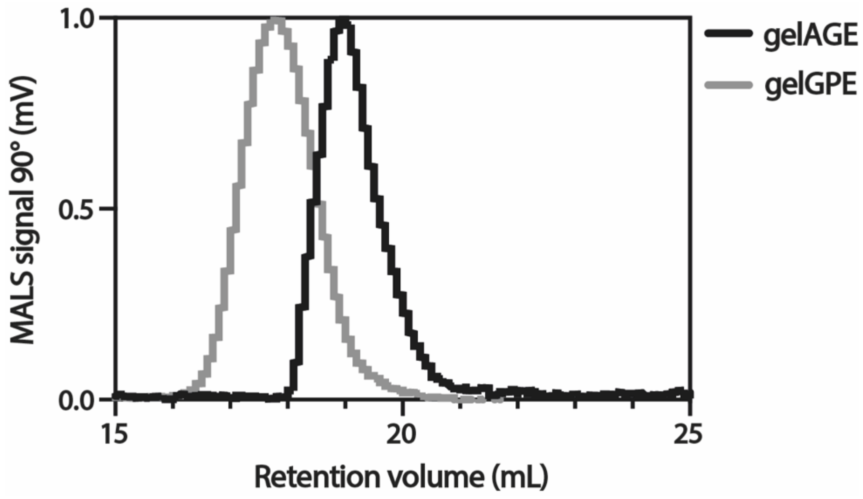


**Supplementary Fig.3: Aqueous-GPC spectra comparison of gelAGE and gelGPE.** Multi angle light scattering (90°-MALS) chromatogram of gelAGE ($G_{1MM}$) in comparison to gelGPE.

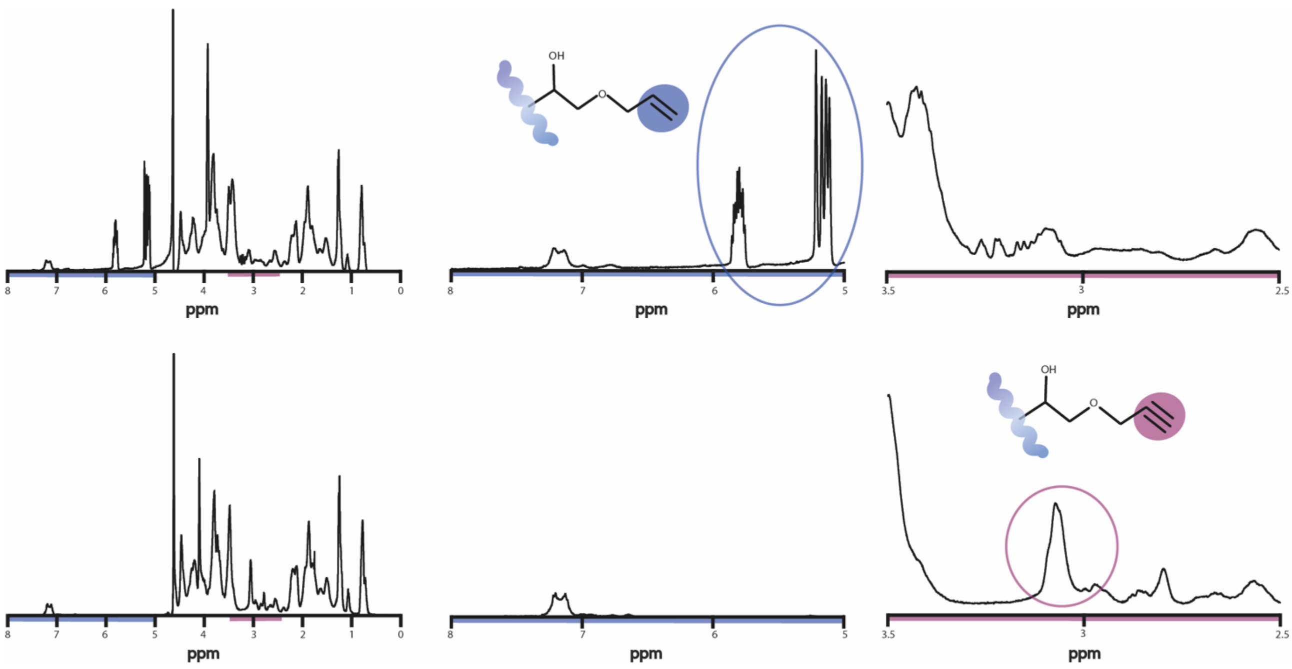


**Supplementary Fig.4: $^{1}$H-NMR spectra comparison of gelAGE and gelGPE.** $^{1}$H-NMR spectrum of a gelAGE (top) and gelGPE sample (bottom). The blue highlighted area of the spectra (δ = 7.8 – 5.0 ppm) contains the phenylalanine peak (δ = 7.45—7.25 ppm) and its integral is normalized for its 5 protons. Two other peaks (δ = 6.0—5.0 ppm) are representing the allyl protons of the gelAGE. The integral value of the single proton (δ = 6.02 – 5.88 ppm) is used to determine the DoM of the gelAGE. The peak centered around δ = 3.15—3.05 ppm refers to the carbon-carbon triple bond grafted on the gelGPE, and its integral is used to determine the DoM of the gelGPE.
