## Supplementary material for "Optical Fiber-Assisted Printing: A Platform Technology for Straightforward Photopolymer Resins Patterning and Freeform 3D Printing": Description of supplementary information files

**Description of Additional Supplementary Files**

File name: Supplementary movie 1
Description: OFAP 2D-photopatterning of a gelAGE-based resin (feed rate: 1 mm $s^{-1}$, gap: 0.15 mm)

File name: Supplementary movie 2
Description: OFAP 3D-embedded printing of a gelAGE-based resin (dye: tartrazine, polymer optical fiber featuring a 500 μm diameter)
